# Cholesterol metabolism and intrabacterial potassium homeostasis are intrinsically related in *Mycobacterium tuberculosis*

**DOI:** 10.1101/2024.11.10.622811

**Authors:** Yue Chen, Berge Hagopian, Shumin Tan

## Abstract

Potassium (K^+^) is the most abundant intracellular cation, but much remains unknown regarding how K^+^ homeostasis is integrated with other key bacterial biology aspects. Here, we show that K^+^ homeostasis disruption (CeoBC K^+^ uptake system deletion) impedes *Mycobacterium tuberculosis* (Mtb) response to, and growth in, cholesterol, a critical carbon source during infection, with K^+^ augmenting activity of the Mtb ATPase MceG that is vital for bacterial cholesterol import. Reciprocally, cholesterol directly binds to CeoB, modulating its function, with a residue critical for this interaction identified. Finally, cholesterol binding-deficient CeoB mutant Mtb are attenuated for growth in lipid-rich foamy macrophages and *in vivo* colonization. Our findings raise the concept of a role for cholesterol as a key co-factor, beyond its role as a carbon source, and illuminate how changes in bacterial intrabacterial K^+^ levels can act as part of the metabolic adaptation critical for bacterial survival and growth in the host.

## INTRODUCTION

Ions are fundamental to cellular physiology^1–6^, and in the context of host-pathogen interactions, extensive research has focused on metal ions such as iron, zinc, and manganese, due to their scarcity and the competition between host and pathogen for their acquisition^3,7,8^. However, abundant ions, such as potassium (K^+^) and chloride (Cl^-^), also play critical roles in host-pathogen interactions, both in driving bacterial transcriptional responses and adaptation, and in their roles in cellular homeostasis^6,9–15^. Of pertinence here, K^+^ is the most abundant intracellular cation in both host and bacterial cells, and its levels must be carefully regulated for proper cellular function^15,16^. In the host, K^+^ plays myriad functions, including as a signal in the induction of immune responses, with for example low K^+^ concentrations ([K^+^]) triggering activation of the NLRP3 inflammasome^17,18^. In bacteria, beyond its most often studied role in osmoprotection^15,19^, disruption of K^+^ homeostasis has also been reported to affect aspects ranging from *Salmonella* effector protein secretion^20^, to *Streptococcus mutans* acid stress adaptation^21^. These studies highlight the critical role of K^+^ in bacterial biology and pathogenicity, but much remains unknown regarding how intrabacterial [K^+^] may be regulated in response to other environmental cues, and the underlying mechanisms that account for the impact of K^+^ homeostasis disruption on bacterial biology phenotypes.

*Mycobacterium tuberculosis* (Mtb), the causative agent of tuberculosis, is a bacterial pathogen highly adapted for colonization of the human host, and remains the leading cause of death from an infectious disease worldwide^22^. The ability of Mtb to respond to ionic signals and maintain intrabacterial ionic homeostasis is critical for its survival within the host. For example, Mtb is adept at maintaining intrabacterial pH near neutrality even in the presence of acidic environmental pH levels, with disruption of this ability resulting in attenuated host colonization^23^. In the case of K^+^, we have previously showed that disruption of Mtb K^+^ homeostasis by deletion of *ceoBC*, encoding the constitutive, low-medium affinity Trk K^+^ uptake system^24^, significantly impaired Mtb response to acidic pH and high [Cl^-^] in its local environment, without affecting intrabacterial pH or membrane potential^13^. Δ*ceoBC* Mtb is consequently attenuated for host colonization in both macrophage and murine infection models^13^, underscoring the importance of K^+^ homeostasis in the biology of Mtb-host interactions.

Crucially, host colonization by bacterial pathogens entails not just adaptation to changing ionic signals, but also integration of these responses to the availability of different nutrient sources during infection. Intriguingly, we recently discovered that a reduction in environmental [K^+^] dampened the transcriptional response of Mtb to cholesterol, while the presence of cholesterol conversely increased induction of K^+^ regulon genes^25^. Lipids, including cholesterol, are a vital carbon source for Mtb during infection, and deletion of Mtb genes needed for cholesterol utilization results in significantly attenuated host colonization^26,27^. How Mtb K^+^ homeostasis might impact bacterial cholesterol metabolism and vice versa remain open questions.

Here, we interrogate this interplay between Mtb K^+^ homeostasis and cholesterol uptake and metabolism. Our work reveals that disruption of K^+^ homeostasis via deletion of the CeoBC Trk K^+^ uptake system impedes Mtb response to, and growth in, cholesterol. This impairment likely arises from decreased intrabacterial [K^+^] in Δ*ceoBC* Mtb diminishing the activity of the ATPase MceG, which is vital for Mtb import of cholesterol^28–30^. Reciprocally, we find that cholesterol directly binds to CeoB, modulating its function, and identify a residue critical for this interaction. The interplay between Mtb K^+^ homeostasis and cholesterol uptake and metabolism is vital for the bacterium’s virulence, as disruption of the cholesterol-binding ability of CeoB results in significant attenuation of Mtb growth in lipid-rich foamy macrophages, and in a murine infection model that recapitulates canonical necrotic granulomas observed during human disease. Our findings raise the concept of a role for cholesterol as a key co-factor, beyond its role as a carbon source, and illuminate how changes in Mtb intrabacterial K^+^ levels act as part of the metabolic adaptation critical for Mtb survival and growth in the host.

## RESULTS

### Disruption of K^+^ homeostasis inhibits Mtb cholesterol response

To examine how K^+^ homeostasis affects Mtb cholesterol response, we tested cholesterol regulon gene expression levels upon exposure of Δ*ceoBC* Mtb to cholesterol. Intriguingly, deletion of *ceoBC* resulted in reduced induction of cholesterol regulon genes as compared to WT Mtb, which was restored upon complementation (*ceoBC*\*) (Figure 1A). In contrast, disruption of the high affinity Kdp K^+^ uptake system, which is induced only in the presence of limiting K^+^, had no effect on the bacterium’s cholesterol response (Figure S1A). In accord with the dampening of the cholesterol transcriptional response, growth of Δ*ceoBC* Mtb was significantly reduced in cholesterol medium, but not in standard 7H9 rich medium (glucose and glycerol as carbon sources; Figure 1B). We next tested if Δ*ceoBC* Mtb were altered in their ability to import cholesterol, utilizing assays with an intrinsically fluorescent cholesterol analog, dehydroergosterol (DHE). DHE fluorescence is limited in the aqueous phase but increases upon binding to cholesterol-binding proteins^31–33^, and has been effectively used previously to demonstrate cholesterol binding to bacterial proteins^34,35^. As expected, in 7H9 medium, DHE signal was low and not different in WT, Δ*ceoBC* or *ceoBC*\* Mtb (Figure 1C). In contrast, DHE signal was significantly higher in cholesterol medium, reflecting uptake of DHE into Mtb (Figure 1C). Notably, there was reduced DHE signal in Δ*ceoBC* Mtb as compared to WT and *ceoBC*\* Mtb, indicating lower levels of DHE uptake into the mutant bacteria (Figure 1C).

**Figure 1.**
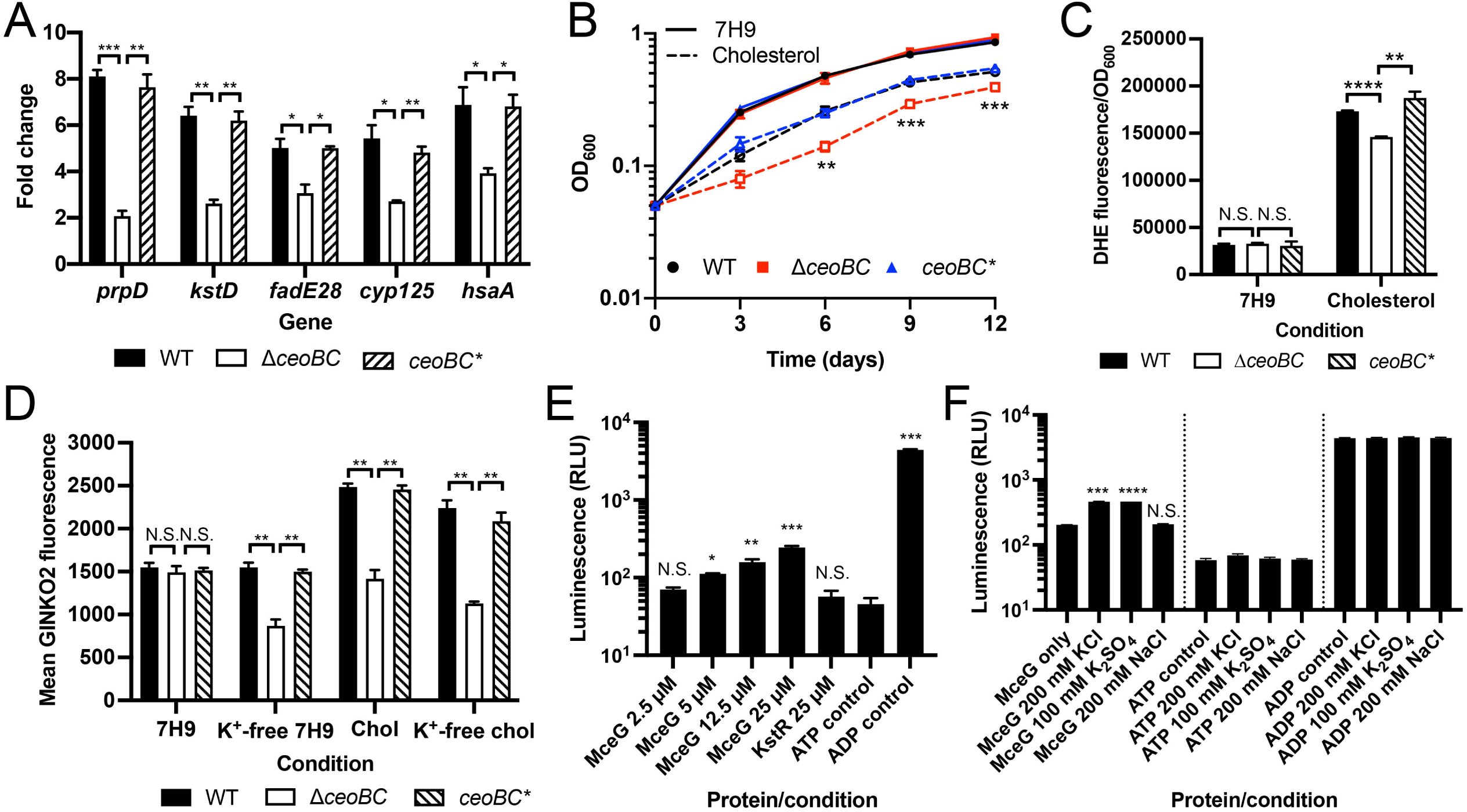
Disruption of K^+^ homeostasis inhibits Mtb cholesterol response. (A) Mtb response to cholesterol is dampened in Δ*ceoBC* Mtb. Log-phase WT, Δ*ceoBC*, and *ceoBC\** (complemented mutant) Mtb were exposed to 7H9 or cholesterol media for 4 hours, before RNA extraction for qRT-PCR analysis. Fold change is as compared to the 7H9 condition, with *sigA* as the control gene. (B) Δ*ceoBC* Mtb is attenuated for growth in cholesterol medium. WT, Δ*ceoBC*, and *ceoBC\** Mtb were grown in 7H9 or cholesterol media, and OD_600_ monitored over time. (C) Cholesterol uptake is reduced in Δ*ceoBC* Mtb. Log-phase WT, Δ*ceoBC*, and *ceoBC\** Mtb were exposed to 7H9 or cholesterol media, supplemented with 100 µM dehydroergosterol (DHE), for 24 hours. DHE uptake into Mtb was measured via analysis of DHE fluorescence on a microplate reader, normalized against OD_600_. (D) Cholesterol increases intrabacterial K^+^ levels in a CeoBC-dependent manner. WT, Δ*ceoBC*, and *ceoBC\** Mtb, each carrying the P_606_’::GINKO2 reporter, were subcultured to OD_600_ = 0.3 into the indicated media (“chol” = cholesterol), and GINKO2 fluorescence measured by flow cytometry 6 days post-assay start. (E) MceG exhibits ATPase activity. His-tagged MceG at indicated concentrations was tested for ATPase activity using an ADP-Glo kinase assay kit. His-tagged KstR1, a transcription factor with no ATPase activity, was also tested. ATP and ADP controls served as negative and positive controls, respectively. Luminescence (relative light units, RLU) was read on a microplate reader. (F) K^+^ increases MceG ATPase activity. 25 µM MceG was incubated with 200 mM KCl, 100 mM K_2_SO_4_ or 200 mM NaCl and tested for ATPase activity as in (E). All data are shown as means ± SEM from three independent experiments. Statistical analyses were performed using an unpaired t-test with Welch’s correction and Holm-Sidak multiple comparisons for (A) – (D). For (B), comparisons were of Δ*ceoBC* to WT in the cholesterol condition. An unpaired t-test with Welch’s correction was used in (E) and (F), with comparisons to the ATP control in (E) and to the no additive control within each group (MceG, ATP, or ADP) for (F). No significance was found for any comparisons in the ATP or ADP control sets in (F). N.S. not significant, * p<0.05, ** p<0.01, *** p<0.001, **** p<0.0001.

To test if cholesterol reciprocally affects K^+^ homeostasis in Mtb, we adapted the genetically encoded GINKO2 K^+^ sensor, which consists of a circularly permuted enhanced GFP integrated with the *Escherichia coli* K^+^ binding protein Kbp^36^, for expression in Mtb. Binding of K^+^ to GINKO2 triggers a conformational change, resulting in a K^+^ concentration-dependent increase in GFP fluorescence^36^. Utilizing Mtb strains constitutively expressing the GINKO2 reporter, we observed that WT and *ceoBC*\* Mtb effectively maintained their intrabacterial K^+^ levels, even after exposure to K^+^-free medium for 6 days (Figure 1D). As expected, Δ*ceoBC* Mtb exhibited significantly reduced GINKO2 reporter signal after growth in K^+^-free conditions, demonstrating disrupted K^+^ homeostasis in the mutant strain (Figure 1D). Interestingly, cholesterol significantly increased intrabacterial [K^+^] in WT and *ceoBC*\* Mtb, a phenotype that was lost in the Δ*ceoBC* mutant (Figure 1D). This was similarly observed in the K^+^-free cholesterol condition (Figure 1D). Disruption of the inducible Kdp high affinity K^+^ uptake system did not affect the increase in intrabacterial [K^+^] in cholesterol medium (Figure S1B), reinforcing that the relationship between cholesterol and K^+^ is specific to basal K^+^ homeostasis and the CeoBC K^+^ uptake system.

K^+^ can serve key roles in enzyme activation^37^, and has been found to be important in activity of ATPases present in systems ranging from archaea to mammalian^37–39^. Markedly, MceG is a Mtb ATPase critical for driving the import of fatty acids and cholesterol through the Mce1 and Mce4 transporters, respectively^28–30^. Given our results above, we thus hypothesized that increased intrabacterial [K^+^] levels stimulate the activity of MceG during Mtb growth in cholesterol medium, enabling the uptake/utilization of cholesterol. As expected, purified MceG exhibited ATPase activity, while a control protein, the transcription factor KstR1 that is involved in cholesterol regulon gene expression control^27,40^, did not (Figure 1E). Intrabacterial [K^+^] has been reported to be in the range of hundreds of millimolar^41^, and strikingly, we found that the presence of increased [K^+^] indeed resulted in higher MceG ATPase activity (Figure 1F). This phenotype was specific to K^+^, with sodium having no effect on MceG ATPase activity (Figure 1F).

Collectively, these data demonstrate that disruption of K^+^ homeostasis impedes the uptake and response of Mtb to cholesterol, consequently decreasing the ability of the bacteria to grow in cholesterol medium. Mechanistically, our findings further indicate that the attenuation of Mtb growth on cholesterol medium upon disruption of intrabacterial K^+^ homeostasis results at least in part from the inability of the mutant Mtb to raise the intrabacterial [K^+^] setpoint in the presence of cholesterol, as K^+^ acts to boost the activity of the MceG ATPase that is crucial in function of the Mce4 cholesterol uptake system.

### Cholesterol directly acts on the CeoBC K^+^ uptake system

Intriguingly, cholesterol has been shown to directly bind to and modulate the activity of the Kir family of K^+^ uptake systems in mammalian cells and the *E. coli* inwardly-rectifying K^+^ channel KirBac1.1^42–45^. Additionally, studies have demonstrated that cholesterol can regulate the function of voltage-gated potassium channels (Kv) in various mammalian systems, including alveolar epithelial and lung cells^46–50^. We thus investigated whether cholesterol directly interacts with CeoBC by employing a thermostability shift assay, which has previously been successfully used to show Mtb protein binding to other factors, such as glycerol and magnesium^51,52^. As shown in Figure 2A, the presence of cholesterol resulted in an increase in CeoB thermostability, which was not observed in the presence of glycerol or glucose. In contrast, no thermostability shift was observed with CeoC in the presence of cholesterol (Figure 2B), demonstrating the specificity of the cholesterol interaction with CeoB. To further verify this interaction, we pursued a second independent approach using DHE. As previously described, DHE fluorescence increases upon binding to a cholesterol-binding protein^31–33^. In accord with the thermostability shift assay, incubation of CeoB with DHE resulted in significantly higher DHE fluorescence signal than with CeoC or the buffer only control (Figure 2C), supporting the specific interaction between cholesterol and CeoB.

**Figure 2.**
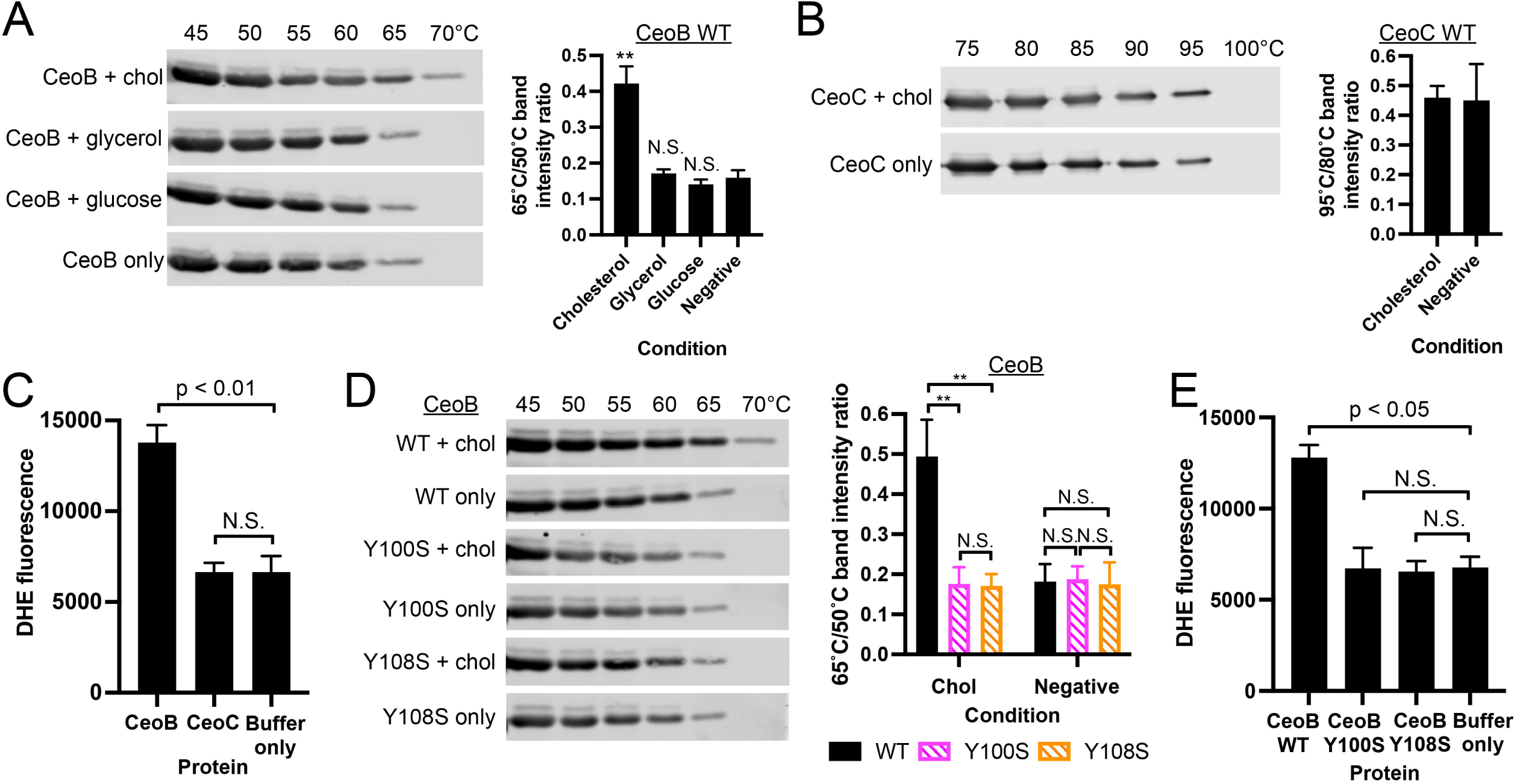
Cholesterol directly acts on the CeoBC K^+^ uptake system. (A and B) CeoB, but not CeoC, shows increased thermostability in the presence of cholesterol. Purified CeoB (A) or CeoC (B) were incubated with 5 µM cholesterol, glycerol, or glucose as noted, at room temperature for 20 minutes, before exposure to indicated temperatures for 5 minutes. Samples were then centrifuged and supernatant aliquots run on SDS-PAGE gels and analyzed by Western blot. Graphs show quantification of band intensity at 65°C/50°C for CeoB (A), or 95°C/80°C for CeoC (B). (C) CeoB, but not CeoC, binds to the fluorescent cholesterol analog dehydroergosterol (DHE). Purified CeoB or CeoC were incubated with 1 µM DHE for 30 minutes, and DHE fluorescence measured on a microplate reader. (D) Y100 and Y108 residues are important for increased CeoB thermostability in the presence of cholesterol. Purified CeoB, CeoB-Y100S, and CeoB-Y108S were tested for thermostability ± cholesterol as in (A). (E) CeoB Y100S and Y108S point mutants are unable to bind DHE. Purified CeoB, CeoB-Y100S and CeoB-Y108S proteins were tested for DHE binding as in (C). Data are shown as means ± SEM from 3-4 experiments for all graphs. p-values were obtained with unpaired t-tests for (A) and (B), a one-way ANOVA (Brown-Forsythe and Welch) with Dunnett’s T3 multiple comparisons test for (C) and (E), and a two-way ANOVA with Tukey’s multiple comparisons test for (D). N.S. not significant, ** p<0.01.

Previous reports, primarily in the context of mammalian cells, have shown cholesterol interaction with proteins at a motif called the “cholesterol recognition amino acid consensus” (“CRAC”) motif (L/V-X_1-5_-Y-X_1-5_-K/R) or the inverted “CARC” version (K/R-X_1-5_-Y-X_1-5_-L/V)^53–55^. Intriguingly, the CeoB sequence contains one CRAC and one CARC motif, which CeoC lacks. To determine whether mutations at these tyrosine (Y) sites disrupt cholesterol binding, we mutated the key Y residues (Y100 for the CRAC motif and Y108 for the CARC motif) to serine (S). Thermostability shift assays showed that both the CeoB Y100S and Y108S mutants lost the ability to bind cholesterol (Figure 2D). Similarly, DHE failed to bind effectively to the CeoB Y100S and Y108S mutant proteins, with DHE fluorescence unchanged from the buffer only control (Figure 2E).

Together, these data demonstrate that cholesterol binds directly to CeoB, part of the CeoBC Trk K^+^ uptake system that is critical for K^+^ homeostasis in Mtb. It further identifies CRAC and CARC motifs, and the Y100 and Y108 residues, as potential key sites of cholesterol interaction with CeoB.

### Cholesterol binding to CeoB is critical for Mtb response and adaptation to cholesterol

Having established that CeoB is able to directly bind to cholesterol, we next sought to determine the biological consequences of this binding to CeoBC function and Mtb biology. We had previously demonstrated that deleting *ceoBC* leads to a significant reduction in Mtb response to acidic pH and high [Cl^-^]^13^. To test the impact of loss of CeoB cholesterol binding on this response, we introduced the pH/Cl-responsive reporter *rv2390c’*::GFP into Δ*ceoBC* complemented with *ceoBC* alleles where CeoB contained either the Y100S or Y108S point mutations (*ceoB(Y100S)C\** or *ceoB(Y108S)C\**, respectively). As shown in Figure 3A, the mutations at Y100 and Y108 did not affect *rv2390c’*::GFP reporter signal induction under high [Cl^-^] conditions. This result supports the conclusion that these mutations specifically disrupt the cholesterol binding of CeoB, and do not affect its function in contexts absent of cholesterol.

**Figure 3.**
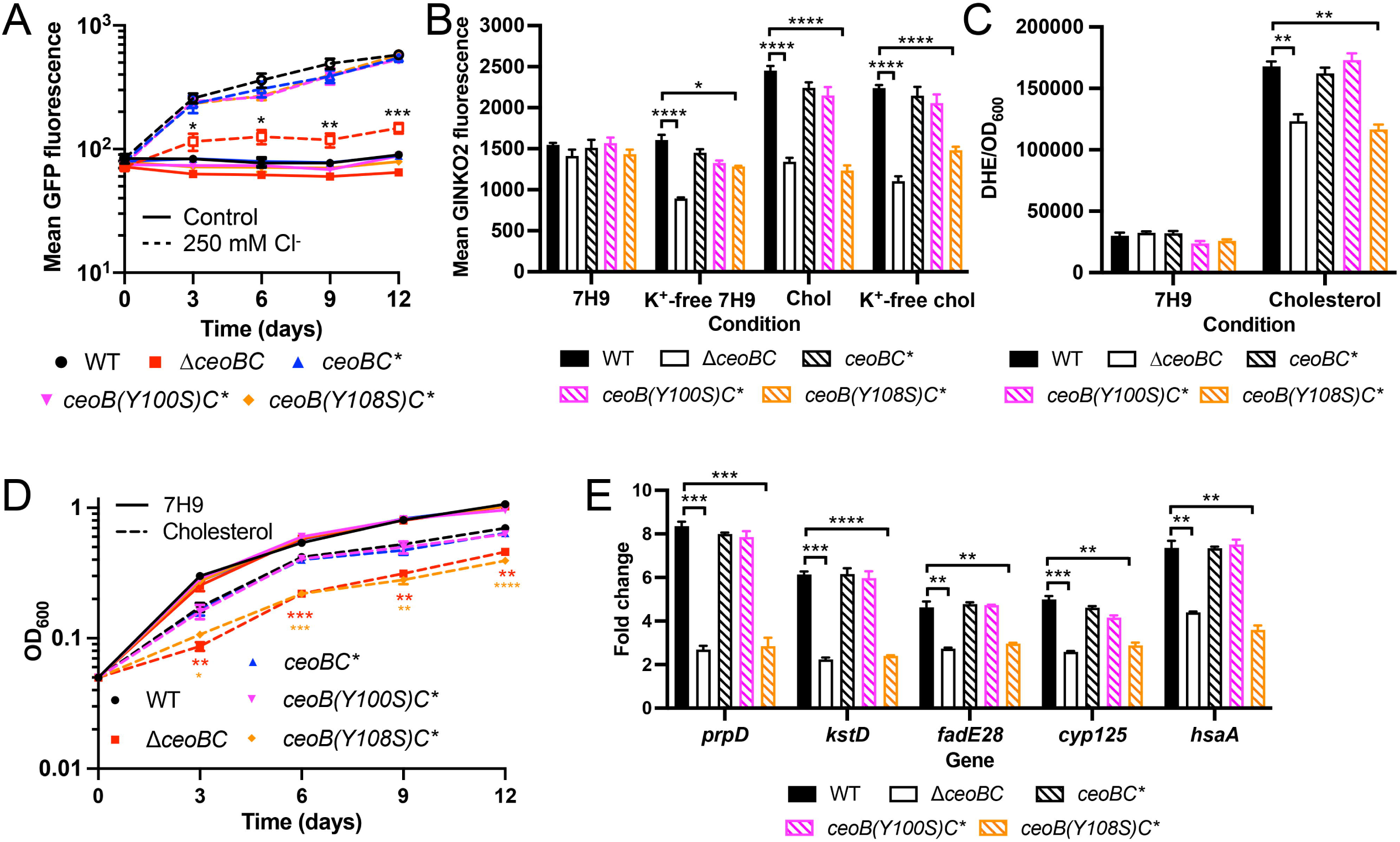
Cholesterol binding to CeoB is critical for Mtb response and adaptation to cholesterol. (A) The ability of CeoB to bind to cholesterol does not affect Mtb response to Cl^-^. WT, *ΔceoBC*, *ceoBC\**, *ceoB(Y100S)C\** and *ceoB(Y108S)C\** Mtb each carrying the Cl^-^-responsive *rv2390c*’::GFP reporter were grown in 7H9 medium ± 250 mM NaCl. Samples were taken at indicated time points and fixed for analysis of reporter expression by flow cytometry. (B) *ceoB(Y108S)C\** Mtb fails to exhibit increased intrabacterial [K^+^] in the presence of cholesterol. WT, *ΔceoBC*, *ceoBC\**, *ceoB(Y100S)C\** and *ceoB(Y108S)C\** Mtb each carrying the P_606_’::GINKO2 reporter were subcultured at OD_600_ = 0.3 into the indicated media (“chol” = cholesterol), and GINKO2 fluorescence measured by flow cytometry 6 days post-assay start. (C) Cholesterol uptake is reduced in *ceoB(Y108S)C\** Mtb. Indicated Mtb strains were exposed to 7H9 or cholesterol media, supplemented with 100 µM DHE, for 24 hours. DHE uptake into Mtb was measured via analysis of DHE fluorescence on a microplate reader, normalized against OD_600_. (D) *ceoB(Y108S)C\** Mtb is attenuated for growth in cholesterol. Indicated Mtb strains were cultured in 7H9 or cholesterol media and growth tracked by OD_600_ over time. (E) *ceoB(Y108S)C\** Mtb has a dampened response to cholesterol. Indicated Mtb strains were exposed to 7H9 or cholesterol media for 4 hours, before RNA extraction for qRT-PCR analysis. Fold change is as compared to the 7H9 condition, with *sigA* as the control gene. Data in all panels are shown as means ± SEM from 3 experiments. p-values were obtained with an unpaired t-test with Welch’s correction and Holm-Sidak multiple comparisons. For (A), comparisons were of the mutant/complement strains to WT in the 250 mM Cl^-^ condition. For (B), (C), and (E), comparisons made were for the mutant/complement strains to WT for each condition. For (D), comparisons were of the mutant/complement strains to WT in the cholesterol condition. Only comparisons with significant p-values are indicated. * p<0.05, ** p<0.01, *** p<0.001, **** p<0.0001.

In agreement with the continued functionality of the CeoB Y100S and Y108S proteins in non-cholesterol-related contexts, the point mutants also did not impair the ability of Mtb to maintain intrabacterial [K^+^] under K^+^-limiting conditions (Figure 3B). However, unlike WT and *ceoBC*\* Mtb, the *ceoB(Y108S)C\** strain exhibited a phenotype similar to Δ*ceoBC* in the presence of cholesterol, failing to increase its intrabacterial [K^+^] (Figure 3B). Surprisingly, despite the purified protein assays indicating that CeoB Y100S is also unable to bind to cholesterol, *ceoB(Y100S)C\** Mtb showed similar increased intrabacterial [K^+^] as WT and *ceoBC*\* in cholesterol-containing conditions (Figure 3B). This finding suggests that the ability of the CeoB Y100S protein to bind cholesterol might be rescued in the context of intact bacteria, where other protein partners, such as CeoC, are present and may affect overall structure of the CeoBC complex. Similarly, cholesterol uptake as indicated by DHE fluorescence showed that only the *ceoB(Y108S)C\** mutant phenocopied Δ*ceoBC*, with decreased DHE signal versus WT when the bacteria were grown in cholesterol medium (Figure 3C).

Examination of the effect of the CeoB point mutations on Mtb growth in, and response to, cholesterol medium reinforced the importance of the Y108 residue for proper cholesterol interaction with CeoB. In particular, the *ceoB(Y108S)C\** mutant exhibited a growth defect identical to Δ*ceoBC* Mtb in cholesterol medium (Figure 3D), with a corresponding reduction in its cholesterol transcriptional response (Figure 3E).

These data identify the Y108 residue as vital for the interaction of cholesterol with CeoB, and demonstrate the critical importance of CeoB binding to cholesterol for proper adaptation of Mtb to cholesterol.

### Cholesterol binding to CeoB is important for Mtb host infection

Levels of cholesterol experienced by Mtb during host infection vary spatiotemporally, with lipid-rich foamy macrophages observed ringing the center necrotic lesions that form as infection progresses, but not at earlier time points^25,56^. We had previously shown that deletion of *ceoBC* resulted in attenuation for Mtb growth in murine bone marrow-derived macrophages (BMDMs)^13^. To test how the presence of lipids may further alter the ability of Δ*ceoBC* to colonize host macrophages, we infected untreated or oleate-treated BMDMs (to induce formation of foamy macrophages^25,57,58^) with WT, Δ*ceoBC*, and the various *ceoBC* complementation strains. Strikingly, we found that Δ*ceoBC* Mtb exhibited even greater attenuation in foamy macrophages compared to untreated BMDMs (Figure 4A). As expected, *ceoB(Y108S)C\** Mtb, but not *ceoB(Y100S)C\** Mtb, phenocopied Δ*ceoBC* Mtb in exhibiting reduced growth both in untreated and foamy BMDMs (Figure 4A).

**Figure 4.**
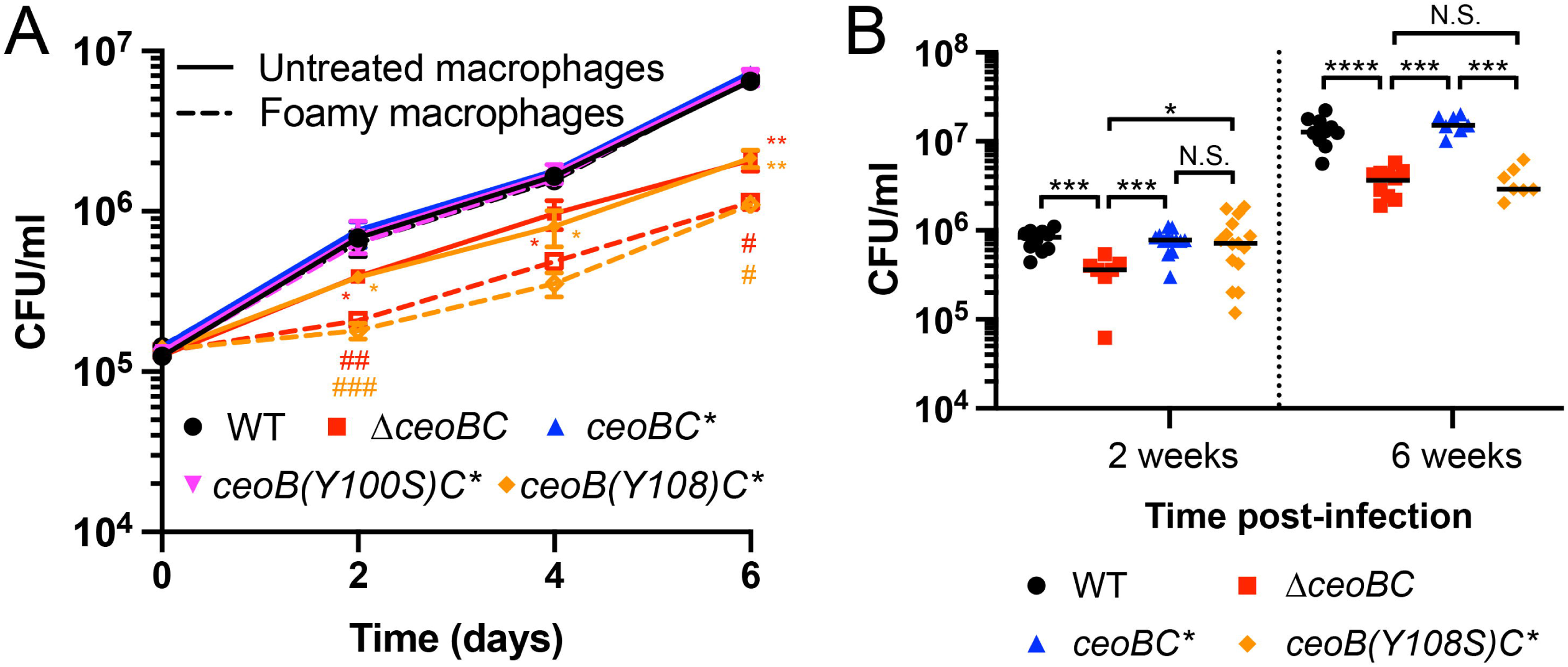
The ability of CeoB to bind to cholesterol is important for host colonization. (A) Δ*ceoBC* and *ceoB(Y108S)C\** Mtb exhibit increased attenuation for growth in foamy versus untreated macrophages. Murine bone marrow-derived macrophages untreated or pre-treated with oleate for 24 hours to induce foamy macrophages were infected with the indicated Mtb strains and colony forming units (CFUs) tracked over time. Data are shown as means ± SD from 3 wells, representative of 3 independent experiments. p-values comparing each strain to WT in the untreated macrophages were obtained with a 2-way ANOVA with Tukey’s multiple comparisons test (*). Unpaired t-tests with Welch’s correction were further applied to compare Δ*ceoBC* and *ceoB(Y108S)C\** infections in foamy versus untreated macrophages (#). Only comparisons with significant p-values are indicated. *, # p<0.05, **, ## p<0.01, ### p<0.001. (B) *ceoB(Y108S)C\** Mtb is attenuated for colonization in a murine infection model when foamy macrophages are present. C3HeB/FeJ mice were infected with the indicated Mtb strains, and lung homogenates plated for CFUs 2 or 6 weeks post-infection. p-values were obtained with a Mann-Whitney statistical test. N.S. not significant, * p<0.05, *** p<0.001, **** p<0.0001.

Finally, to assess the role of cholesterol binding to CeoB in the context of Mtb infection of a whole animal host, C3HeB/FeJ mice were infected with WT, *ΔceoBC*, *ceoBC\**, or *ceoB(Y108S)C\** Mtb and bacterial loads determined at 2 and 6 weeks post-infection. Hallmark necrotic lesions are formed in this mouse strain upon Mtb infection, with lipid-rich foamy macrophages present at 6 weeks, but not 2 weeks, post-infection^25^. At both time points examined, a significant reduction in bacterial load was observed for the *ΔceoBC* mutant compared to WT (Figure 4B), consistent with our previous findings of attenuation in host colonization of Δ*ceoBC* Mtb in the C57BL/6J murine infection model^13^. In contrast, a defect in host colonization of *ceoB(Y108S)C\** Mtb was observed only at 6 weeks, but not 2 weeks, post-infection (Figure 4B). Together, these data demonstrate the importance of cholesterol binding to CeoB for Mtb growth and survival in lipid-rich environments during host infection.

## DISCUSSION

While K^+^ has been well-appreciated as the most abundant cation present in bacterial cells^15^, how maintenance of K^+^ homeostasis may relate to bacterial metabolism adaptation, and how K^+^ homeostasis itself is regulated in response to changing nutrient conditions, has remained open questions. Our findings here reveal the critical interplay between K^+^ homeostasis and cholesterol metabolism in Mtb, and support two concepts that are likely to have broad pertinence across bacterial species, given the fundamental role of K^+^ in bacterial biology.

First, the observation that the intrabacterial [K^+^] setpoint is increased in the presence of cholesterol, with K^+^ acting to stimulate the activity of the ATPase MceG needed for cholesterol uptake and utilization by Mtb, supports the concept that K^+^ homeostasis is dynamic and integrated with environmental signals, with K^+^ serving a role in the regulation of key downstream pathways. Ions as critical cofactors for enzymes is well-appreciated in all kingdoms, with scarce divalent cations such as iron, manganese, and zinc being the most intensely studied^59,60^. In mammalian and plant biology however, the important role that K^+^ can play in enzyme activity has also been recognized^37,61^. This includes in kinases such as branched-chain ⍺-ketoacid dehydrogenase and pyruvate dehydrogenase kinase, where K^+^ acts to critically stabilize parts of the protein^62,63^, and in ATPases such as Hsc70 and the plant plasma membrane H^+^-ATPase proton pump, which exhibit higher ATPase activity in the presence of K^+38,64^. Our results here with MceG and the change in intrabacterial [K^+^] in response to changes in external environment, in combination with the status of K^+^ as the most abundant intracellular cation in bacterial cells^15^, suggest that similar dependencies on K^+^ are likely to also exist more widely in the bacterial kingdom. We propose that future studies examining the role of K^+^ will yield vital insight into a new facet of regulation of bacterial enzymatic activities. They will also provide understanding of whether K^+^ acts in concert with other divalent cations such as magnesium or manganese, as is often the case in the K^+^-regulated mammalian enzymes studied to date^37,61^.

Second, our finding that cholesterol directly binds to CeoB, a component of the Trk K^+^ uptake system in Mtb, affecting its function and Mtb host colonization in the context of lipid-rich environments, raises the concept of cholesterol as not just a carbon source, but also a molecule capable of regulating key facets of bacterial biology. Cholesterol binding to K^+^ transport systems in mammalian cells can result in either upregulation or downregulation of K^+^ transport^42,65^. Here, cholesterol binding to CeoB appears to increase activity of the Trk K^+^ uptake system, given the observed increase in intrabacterial K^+^ levels. The one previous example, to our knowledge, of a bacterial K^+^ uptake system affected by cholesterol is KirBac1.1 from *E. coli*, where cholesterol was found to inhibit channel activity^44,45^. Those studies were however conducted with purified proteins incorporated into liposomes, with radioactive rubidium (^86^Rb^+^) as a proxy for K^+^ transport measurement^44,45^. Unlike mammalian K^+^ transport systems, bacterial K^+^ transport systems often discriminate against Rb^+66–68^; our establishment of the GINKO2 reporter for relative measurement of intrabacterial K^+^ in intact bacteria opens the path for future studies examining how cholesterol, or other potential co-factors, affect bacterial K^+^ uptake in physiological context. Further, our identification of the Y108 residue, part of a CARC motif, as essential for cholesterol binding to CeoB and the effects of cholesterol on CeoB function in intact Mtb cells sets the foundation for mechanistic understanding. Future studies could be aimed at unveiling the precise mechanism by which cholesterol binding to CeoB elevates K^+^ levels within Mtb; perhaps, for example, by structural-based changes to the uptake system, as has been identified in mammalian systems^42^.

The CRAC cholesterol recognition motif was originally put forth from a study focused on the mammalian peripheral-type benzodiazepine receptor that regulates the transport of cholesterol across the mitochondrial outer and inner membranes^69^, with later studies describing the inverted CARC motif, and the presence and role of both motifs for various mammalian cholesterol-binding proteins^53,70–72^. In bacterial systems, studies that have examined the CRAC/CARC motifs have largely centered on toxins that interact with cholesterol present in the host cell membrane during the process of target cell intoxication, such as ⍺-hemolysin from *E. coli*, cytolethal distending toxins from *Campylobacteri jejuni* and *Aggregatibacter actinomycetemcomitans*, and leukotoxin from *A. actinomycetemcomitans*^54,73–75^. For bacteria such as Mtb, *Rhodococcus* sp., and *Gordonia* sp. that can utilize cholesterol as a carbon source^26,27,76–79^, the possibility of cholesterol functioning in diverse aspects of its biology are greatly expanded, given the active uptake of cholesterol into the bacterium and the consequent presence of cholesterol within the bacterial cell. Intriguingly, examination of the Mtb genome reveals the presence of CRAC and CARC motifs in a wide variety of proteins, spanning transporters to transcription factors. However, in studies with mammalian systems and bacterial toxins, not all previously identified CRAC and CARC motifs have been found to impact cholesterol binding^43,53,80^, which is perhaps unsurprising given the relatively loose sequence definition of the motifs. We propose that further study of the possible role of cholesterol in the function of Mtb and other bacterial proteins that encode CRAC or CARC motifs will aid in better defining these motifs, and importantly lead to new discoveries regarding cholesterol-driven regulation of bacterial biology.

Ionic homeostasis and environmental and metabolic adaptation are critical facets for all bacteria, and our results here open new avenues in the understanding of how these facets are integrated. Application and extension of the concepts raised here thus hold exciting potential for revealing both fundamental insight into bacterial biology and key integration nodes that can be targeted to disrupt successful adaptation to local niches by a pathogen.

## ACKNOWLEDGEMENTS

We thank members of the Tan laboratory for helpful discussion. This work was supported by grants from the National Institutes of Health (R01 AI143768 and R21 AI171356) to ST, and by an American Lung Association Catalyst Award (CA-1268681) to YC.

## AUTHOR CONTRIBUTIONS

Conceptualization, YC and ST; Investigation, YC, BH, and ST; Writing – original draft, YC and ST; Writing – review and editing, YC, BH, and ST; Supervision, ST; Funding acquisition, YC and ST.

## STAR★Methods

### Key resources table

| REAGENT or RESOURCE | SOURCE | IDENTIFIER |
| --- | --- | --- |
| Antibodies |  |  |
| Mouse anti-6x-His-tag | Invitrogen | R930-25 |
| Mouse anti-FLAG | Sigma | F3165 |
| Goat anti-mouse IRDye 680RD | LI-COR | 926-68070 |
| Bacterial strains |  |  |
| *Mycobacterium tuberculosis* CDC1551 streptomycin-resistant | Lab stock | N/A |
| *Mycobacterium tuberculosis* CDC1551 streptomycin-resistant $\Delta ceoBC$ | MacGilvary et al.<sup>13</sup> | N/A |
| *Mycobacterium tuberculosis* CDC1551 streptomycin-resistant *ceoBC\** (complemented $\Delta ceoBC$ strain) | MacGilvary et al.<sup>13</sup> | N/A |
| *Mycobacterium tuberculosis* CDC1551 streptomycin-resistant *ceoB(Y100S)C\** | This manuscript | N/A |
| *Mycobacterium tuberculosis* CDC1551 streptomycin-resistant *ceoB(Y108S)C\** | This manuscript | N/A |
| *Mycobacterium tuberculosis* CDC1551 streptomycin-resistant *rv2390c*::GFP | Tan et al.<sup>12</sup> | N/A |
| *Mycobacterium tuberculosis* CDC1551 streptomycin-resistant $\Delta ceoBC$, *rv2390c*::GFP | MacGilvary et al.<sup>13</sup> | N/A |
| *Mycobacterium tuberculosis* CDC1551 streptomycin-resistant *ceoBC* * (complemented $\Delta ceoBC$ strain), *rv2390c*::GFP | MacGilvary et al.<sup>13</sup> | N/A |
| *Mycobacterium tuberculosis* CDC1551 streptomycin-resistant *ceoB* (Y100S) *C* *, *rv2390c*::GFP | This manuscript | N/A |
| *Mycobacterium tuberculosis* CDC1551 streptomycin-resistant *ceoB* (Y108S) *C* *, *rv2390c*::GFP | This manuscript | N/A |
| *Mycobacterium tuberculosis* CDC1551 streptomycin-resistant P<sub>606</sub>::GINKO2 | This manuscript | N/A |
| *Mycobacterium tuberculosis* CDC1551 streptomycin-resistant $\Delta ceoBC$, P<sub>606</sub>::GINKO2 | This manuscript | N/A |
| *Mycobacterium tuberculosis* CDC1551 streptomycin-resistant *ceoBC* * (complemented $\Delta ceoBC$ strain), P<sub>606</sub>::GINKO2 | This manuscript | N/A |
| *Mycobacterium tuberculosis* CDC1551 streptomycin-resistant *ceoB* (Y100S) *C* *, P<sub>606</sub>::GINKO2 | This manuscript | N/A |
| *Mycobacterium tuberculosis* CDC1551 streptomycin-resistant *ceoB* (Y108S) *C* *, P<sub>606</sub>::GINKO2 | This manuscript | N/A |
| *Mycobacterium tuberculosis* CDC1551 streptomycin-resistant $\Delta kdpBC$ | This manuscript | N/A |
| *Mycobacterium tuberculosis* CDC1551 streptomycin-resistant *kdpBC* * (complemented $\Delta kdpBC$ strain) | This manuscript | N/A |
| *Mycobacterium tuberculosis* CDC1551 streptomycin-resistant $\Delta kdpBC$, P<sub>606</sub>::GINKO2 | This manuscript | N/A |
| *Mycobacterium tuberculosis* CDC1551 streptomycin-resistant *kdpBC* * (complemented $\Delta kdpBC$ strain), P<sub>606</sub>::GINKO2 | This manuscript | N/A |
| *Mycobacterium tuberculosis* Erdman streptomycin-resistant | Lab stock | N/A |
| *Mycobacterium tuberculosis* Erdman streptomycin-resistant $\Delta ceoBC$ | MacGilvary et al.<sup>13</sup> | N/A |
| *Mycobacterium tuberculosis* Erdman streptomycin-resistant *ceoBC* * (complemented $\Delta ceoBC$ strain) | MacGilvary et al.<sup>13</sup> | N/A |
| *Mycobacterium tuberculosis* Erdman streptomycin-resistant *ceoB* (Y100S) *C* * | This manuscript | N/A |
| *Mycobacterium tuberculosis* Erdman streptomycin-resistant *ceoB* (Y108S) *C* * | This manuscript | N/A |
| Chemicals, peptides, and recombinant proteins |  |  |
| Dehydroergosterol | Sigma | E2634 |
| Critical commercial assays |  |  |
| ADP-Glo kinase assay | Promega | V6930 |
| Experimental models: Cell lines |  |  |
| L-929 | ATCC | CCL-1 |
| Experimental models: Organisms/strains |  |  |
| Mouse: C57BL/6J | Jackson Laboratories | 000664 |
| Mouse: C3HeB/FeJ | Jackson Laboratories | 000658 |
| Oligonucleotides |  |  |
| See Table S1 for list of oligonucleotides used. |  |  |
| Software and algorithms |  |  |
| FlowJo | BD | https://www.flowjo.com |
| Prism | GraphPad Software | https://www.graphpad.com |
| Image Studio | LI-COR | https://www.licor.com/bio/image-studio/ |

### Resource availability

Further information and requests for resources and reagents should be directed to and will be fulfilled by the lead contact, Shumin Tan.

### Materials availability

All newly generated Mtb strains are available on request from the lead contact, to investigators with the necessary biosafety level 3 facilities to receive and work with these materials.

### Data and code availability

- All data reported in this paper will be shared by the lead contact upon request.
- This paper does not report original code.
- Any additional information required to reanalyze the data reported in this work paper is available from the lead contact upon request

### Experimental model and subject details

#### Murine strains

C57BL/6J and C3HeB/FeJ wild type female mice were obtained from Jackson Laboratories and were 4 weeks old on arrival. Mtb infections were carried out when the mice were 6 weeks of age. All animal protocols in this research followed The National Institutes of Health “Guide for Care and Use of Laboratory Animals” guidelines. All animal protocols (#B2024-90) were reviewed and approved by the Institutional Animal Care and Use Committee at Tufts University, in accordance with guidelines from the Association for Assessment and Accreditation of Laboratory Animal Care, the US Department of Agriculture, and the US Public Health Service.

### Method details

#### Mtb strains and culture

Mtb cultures were propagated as previously described^81^, with all media buffered to pH 7 with 100 mM MOPS and antibiotics added to the media as needed at the following concentrations: 100 µg/ml streptomycin, 50 µg/ml hygromycin, 50 µg/ml apramycin, and 25 µg/ml kanamycin. Strains for *in vitro* assays were in the CDC1551 background, and those for *in vivo* assays were in the Erdman background. The *rv2390c’*::GFP, Δ*ceoBC* mutant and its complement strain, with and without the *rv2390c’*::GFP reporter, have all been previously described^13^. The Δ*kdpBC* mutant and its complement were constructed as previously described^12^, with the Δ*kdpBC* mutation consisting of a deletion beginning at nucleotide 136 of the *kdpB* open reading frame through nucleotide 302 of the *kdpC* open reading frame (as annotated in the Erdman Mtb strain). Complementation of the Δ*kdpBC* mutant was with a construct containing the *kdpBC* operon driven by the *kdpF* promoter, introduced in single copy into the Mtb genome via the pMV306 integrating plasmid. The *ceoB(Y100S)C\** and *ceoB(Y108S)C\** point mutations were constructed using QuikChange mutagenesis (Agilent). The P_606_’::GINKO2 construct was generated by cloning an Mtb codon-optimized GINKO2 (GenScript)^36^, driven by the P_606_ promoter, into the destination Gateway vector pDE43-MEK using the Gateway system^82,83^.

#### qRT-PCR analyses

For qRT-PCR analyses, log-phase Mtb cultures (OD_600_ ∼0.6) were used to inoculate standing T25 flasks with filter caps at an OD_600_ = 0.3, containing 10 ml of 7H9 or cholesterol media. Cholesterol medium with 200 µM cholesterol was prepared as previously described^25,84^. Bacteria were incubated in each medium type for 4 hours, before RNA was extracted as previously described^85^. qRT-PCR experiments were conducted and analyzed according to previously established protocols^13^. Briefly, cDNA was synthesized from 250 ng of extracted RNA using the iScript cDNA synthesis kit (Bio-Rad). qRT-PCR was performed using the iTaq Universal SYBR Green Supermix kit (Bio-Rad) on an Applied Biosystems StepOnePlus real-time PCR system, with each sample run in triplicate. The housekeeping gene *sigA* served as the control, and fold induction was determined using the ΔΔCT method^86^.

#### Dehydroergosterol uptake assays

For dehydroergosterol (DHE; ergosta-5,7,9(11),22-tetraen-3β-ol, Sigma) uptake assay in Mtb, bacteria were cultured to log-phase (OD_600_ ∼0.6) in standing T25 flasks with filter caps in 7H9 medium. Strains were then subcultured to OD_600_ = 0.3 in 7H9 or cholesterol medium, supplemented with 100 µM DHE. 200 μl triplicate aliquots per strain/condition were taken and placed in a clear bottom black 96-well plate (Corning Costar), and incubated for 24 hours. Fluorescence intensity was subsequently measured on a Biotek Synergy Neo2 microplate reader, with excitation 338 nm, emission 381 nm.

#### Recombinant protein expression and purification

To purify CeoB, CeoB-Y100S, and CeoB-Y108S, the genes were cloned into the pET23a plasmid to generate C-terminal 6xHis-tagged proteins. *mceG* (*rv0655*) and *kstR1* (*rv3574*) were each cloned into the pET28a plasmid, generating N-terminal 6xHis-tagged proteins. The CeoC protein was purified by adding a C-terminal Flag tag via PCR and then cloning it into the pET23a vector. Expression plasmids were transformed into *Escherichia coli* BL21(DE3) for recombinant expression and purification. 1 ml of an overnight *E. coli* culture started from frozen stock was used to inoculate 1 L of LB medium + 50 μg/mL ampicillin or kanamycin. Cultures were grown at 37°C, 160 rpm, to an OD_600_ of ∼0.6. Protein production was induced with 1 mM IPTG, and cultures were grown for an additional 4 hours at 37°C, 160 rpm. The supernatants were removed, and cell pellets stored at -80°C prior to further processing.

Purification of the CeoC-FLAG-tagged protein and CeoB, CeoB-Y100S and CeoB-Y108S 6xHis-tagged proteins followed previously described protocols^13,87^. MceG 6xHis-tagged protein was present in the insoluble fraction and was purified via treatment of the insoluble fraction with 5M urea buffer, followed by the standard 6xHis-tagged protein purification protocol^13^. CeoB, CeoB-Y100S, CeoB-Y108S, CeoC, and KstR1 proteins were dialyzed into phosphate buffered saline (PBS) buffer. MceG protein was dialyzed into ATPase reaction buffer (50 mM Tris-HCl, 1 mM MgCl_2_, pH 7.5). Protein concentrations were quantified using a Bradford assay (Bio-Rad).

#### ATPase activity assay

MceG ATPase activity was measured using the ADP-Glo Kinase assay kit (Promega) following the manufacturer’s instruction. Briefly, purified MceG at the indicated concentrations, or 25 µM KstR1, was mixed with 10 µM ATP and ATPase reaction buffer (50 mM Tris-HCl, 1 mM MgCl_2_, pH 7.5) in a total volume of 25 µl and incubated at room temperature for 30 minutes. For the treated groups, indicated concentrations of different compounds (KCl, NaCl, or K_2_SO_4_) were included in the ATPase reaction buffer. After incubation, 25 µl of ADP-Glo reagent was added to each reaction mixture and incubated at room temperature for a further 30 minutes to stop the reaction and deplete unused ATP. Then, 50 µl of kinase detection reagent was added to convert ADP to ATP and introduce luciferin for ATP measurement, followed by a 30 minute incubation in the dark. Triplicate 90 µl aliquots of the final reaction were transferred into a clear-bottom white 96-well plate (Corning Costar) for each sample and luminescence measured with a BioTek H1 multimode microplate reader.

#### Cholesterol binding assays

For thermostability shift cholesterol binding assays, purified proteins (CeoB, CeoB-Y100S, CeoB-Y108S at 15 µg and CeoC at 2.5 µg) were incubated with 5 µM cholesterol, glycerol, or glucose at room temperature for 20 minutes. The mixtures were then incubated at the indicated temperatures for 5 minutes (for CeoB and its point mutants) or 20 minutes (for CeoC). After incubation, the samples were centrifuged at 4°C for 30 minutes and the supernatant then run on 12% SDS-PAGE gels. Gels were transferred to Millipore Immobilon-FL PVDF membranes and blocked overnight at 4°C with LI-COR Odyssey blocking buffer. The membranes were then incubated with either mouse anti-6x-His-tag antibody (Invitrogen) or mouse anti-FLAG antibody (Sigma) as needed, at a dilution of 1:1000 for 1 hour at room temperature. Following incubation, the membranes were washed three times for 5 minutes each with PBS + 0.1% Tween 20. They were then incubated with goat anti-mouse IRDye 680RD (LI-COR) at a dilution of 1:3000 for 1 hour at room temperature. After incubation, the membranes were washed three times for 5 minutes each with PBS + 0.1% Tween 20 and then rinsed with deionized water. Protein bands were visualized using a LI-COR Odyssey CLx imaging system, and signal intensities quantified using Image Studio software (LI-COR).

Protein binding assays with DHE (Sigma) were carried out following a previously published protocol^32,34^. Briefly, purified proteins (10 µM) were mixed with 1 µM DHE in DHE buffer (10 mM HEPES, pH 7.4, and 150 mM NaCl) in a 200 µl volume. The mixture was transferred into a black, clear bottom 96-well plate and incubated at room temperature for 30 minutes in the dark. Fluorescence was measured on a BioTek H1 multimode microplate reader with excitation 338 nm, emission 381 nm.

#### Mtb reporter assays and growth assays

For GINKO2 reporter assays, strains carrying the P_606_’::GINKO2 were propagated to log phase and subcultured to an OD_600_ = 0.3 in (i) 7H9, (ii) K^+^-free 7H9, (iii) cholesterol medium, or (iv) K^+^-free cholesterol medium, in standing T25 flasks with filter caps. Prior to resuspending the cultures in the final assay media, an additional wash step with K^+^-free 7H9 medium was performed. All cultures were then incubated at 37°C for 6 days. After 6 days, aliquots were taken and fixed in 4% paraformaldehyde (PFA) in PBS. The fixed samples were pelleted and then resuspended in PBS + 0.1% Tween 80 for flow cytometry analysis on a BD FACSCalibur. Just prior to running, each sample was passed six times through a tuberculin syringe (25G × 5/8” needle) to disrupt clumps. Reporter signal from 10,000 Mtb cells per sample per experimental run were obtained, with three independent runs conducted. Mean fluorescence value for each sample was determined using FlowJo software (BD).

*rv2390c’*::GFP reporter assays were performed as previously described^12^. In brief, strains were propagated to log phase and subcultured to an OD_600_ = 0.05 in 7H9 medium ± 250 mM NaCl, in standing T25 flasks with filter caps. At each time point, aliquots were taken and fixed in 4% PFA in PBS. Reporter signal was analyzed via flow cytometry as described above.

For growth assays, log-phase Mtb cultures were used to inoculate 10 ml of 7H9 or cholesterol medium at a starting OD_600_ = 0.05 in standing T25 flasks with filter caps. OD_600_ was measured at indicated time points.

#### Macrophage culture and infections

Bone marrow-derived macrophages were isolated from C57BL/6J wild type mice procured from Jackson Laboratories. The cells were cultured in DMEM containing 10% FBS, 10% L929-cell conditioned media, 2 mM L-glutamine, 1 mM sodium pyruvate, and antibiotics (penicillin/streptomycin) as needed. They were maintained in a 37°C incubator with 5% CO_2_. To generate foamy macrophages, cells were pre-treated with macrophage medium supplemented with oleate/albumin complexes (0.42 mM sodium oleate, 0.35% BSA) 24 hours before infection, as previously described^57,58^. Infections of macrophages with Mtb were performed as previously described^12,81^. For colony forming unit (CFU) enumeration, macrophages were lysed in water containing 0.01% sodium dodecyl sulfate (SDS), and serial dilutions plated on 7H10 agar plates containing 100 μg/ml cycloheximide.

#### Mouse Mtb infections

C3HeB/FeJ wild-type mice from Jackson Laboratories were intranasally infected with 10^3^ CFUs of Mtb in a 35 μL volume while under light anesthesia using 2% isoflurane^25,84,88^. At 2 or 6 weeks post-infection, the mice were sacrificed using CO_2_. The left lobe and accessory right lobe of the lungs were then homogenized in PBS + 0.05% Tween 80. Serial dilutions of the lung homogenates were plated on 7H10 agar plates supplemented with 100 μg/ml cycloheximide to quantify CFUs.

#### Quantification and statistical analysis

All statistical analyses were performed using GraphPad Prism. Specific statistical tests conducted are described in the figure legends. A p-value of less than 0.05 was considered statistically significant.

## SUPPORTING INFORMATION FIGURE LEGEND

Figure S1. Disruption of the inducible high affinity Kdp K^+^ transport system does not affect Mtb response to cholesterol or intrabacterial [K^+^]. (A) Deletion of *kdpBC* does not affect Mtb response to cholesterol. WT, *ΔkdpBC*, and *kdpBC\** (complemented mutant) Mtb were exposed to 7H9 or cholesterol medium for 4 hours, before RNA extraction for qRT-PCR analysis. Fold change is as compared to the 7H9 condition, with *sigA* as the control gene. (B) Deletion of *kdpBC* does not affect intrabacterial [K^+^] in Mtb. WT, *ΔkdpBC*, and *kdpBC\** Mtb each carrying the P_606_’::GINKO2 reporter were subcultured to OD_600_ = 0.3 into: (i) 7H9 medium, (ii) K^+^-free 7H9 medium, (iii) cholesterol medium, or (iv) K^+^-free cholesterol medium. GINKO2 fluorescence was measured by flow cytometry 6 days post-assay start. Data in both panels are shown as means ± SEM from 3 experiments. p-values were obtained with an unpaired t-test with Welch’s correction and Holm-Sidak multiple comparisons. N.S. not significant, * p<0.05.

