## Supplementary figures and images for "Cholesterol metabolism and intrabacterial potassium homeostasis are intrinsically related in *Mycobacterium tuberculosis*"

### Figure S1

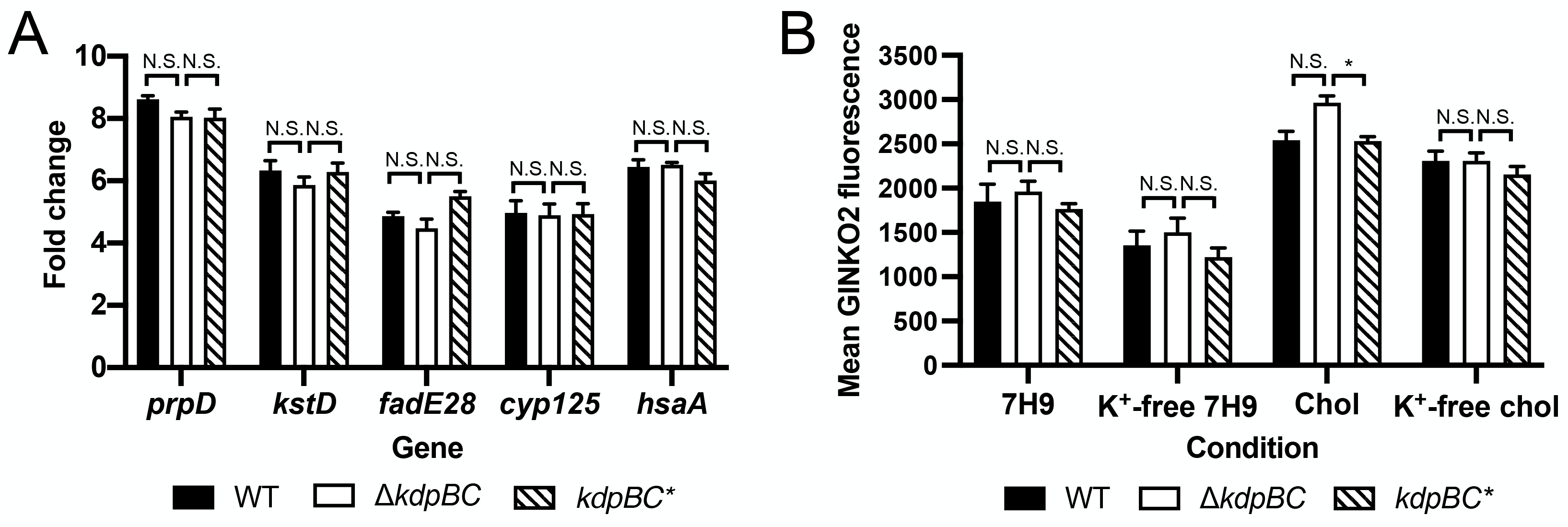
