## Supplementary material for "Cholesterol metabolism and intrabacterial potassium homeostasis are intrinsically related in *Mycobacterium tuberculosis*": Table S1

**Table S1. List of oligonucleotides used.**

| Oligonucleotide | SOURCE | IDENTIFIER |
|---|---|---|
| Primer: GW-GINKO2-F1:<br>5’GGGGACAGCTTTCTTGTACAAAGTGG<br>AGGAGGTATCTCCATGGTGTCGAAGGGCGAG<br>GAG | This manuscript | N/A |
| Primer: GW-GINKO2-R1:<br>5’GGGGACAACTTTGTATAATAAAGTTG<br>TCACATGTACAGCTCGTCCATGCC | This manuscript | N/A |
| Primer: ceoB-Y100S-F:<br>5’TCGTCGCGCGGATCAGTGATGCCAAGCGCG | This manuscript | N/A |
| Primer: ceoB-Y100S-R:<br>5’ CGCGCTTGGCATCACTGATCCGCGCGACG | This manuscript | N/A |
| Primer: ceoB-Y108S-F:<br>5’ GCGCGCCGAGGTCAGTGAGCGACTCGGC | This manuscript | N/A |
| Primer: ceoB-Y108S-R:<br>5’ GCCGAGTCGCTCACTGACCTCGGCGCGC | This manuscript | N/A |
| Primer: ceoB-XbaI-F1:<br>5’ GCTAGG tctaga GAG AGC CAA GGG TAG<br>GCC TAT CCG A | This manuscript | N/A |
| Primer: ceoB-nostop-HindIII-R1:<br>5’ GCTAGG aagctt TCG TCG AGC CCC CGA CTC<br>GAA | This manuscript | N/A |
| Primer: ceoC-BamHI-F1:<br>5’ GCTACC ggatcc ATG AAA GTA GCT GTC GCC<br>GGA GCG | This manuscript | N/A |
| Primer: ceoC-FLAG-HindIII-R2:<br>5’ GCTAGG aagctt TCA CTT GTC GTC GTC ATC<br>CTT GTA GTC CAC GGC CTT GCC GGC AAA<br>CGA GGC GGC CGA CAT GGA CGG CAG CAG<br>CAG CCT | This manuscript | N/A |
| Primer: mceG-nostart-NdeI-F2:<br>5’ GCTAGG catatg GGC GTC AGC ATC GAG GTC<br>AAC | This manuscript | N/A |
| Primer: mceG-BamHI-R1:<br>5’ GCTACC ggatcc TTA CTG GCC GAT TTC GTG<br>CAC | This manuscript | N/A |
| Primer: kdpBC-KO-F1:<br>5’ GCTAGG cttaag GGC CAT CAT GGG CAT CAT<br>TTG GAT CGG AAT GTC A | This manuscript | N/A |
| Primer: kdpBC-KO-R1:<br>5’ GCTAGG tctaga CCT GTA CCC GCG GAG CCA<br>GCT T | This manuscript | N/A |

| | | |
|---|---|---|
| Primer: kdpBC-KO-F2:<br>5’ GCTAGG aagctt GGT CAC CGC CTA CCG CAA GGA A | This manuscript | N/A |
| Primer: kdpBC-KO-R2:<br>5’ GCTAGG actagt GCG CAT CGC GTC GAT TCC GAT CT | This manuscript | N/A |
| Primer: kdpBC-seq-F1:<br>5’ GGC GGG TTC TTC AAC GTG AAC TC | This manuscript | N/A |
| Primer: kdpBC-seq-R1:<br>5’ GAG GTC GCC ACC ATG CCA CTT | This manuscript | N/A |
| Primer: kdpFp-XbaI-F1:<br>5’ GCTAGG tctaga CGG ACG ATC TCG TCG GGG ATC TTC TCC TTC TGC TCG | This manuscript | N/A |
| Primer: kdpFp-NdeI-R1:<br>5’ GCTAGG catatg TCG GCC GGC GCT CGC GTG A | This manuscript | N/A |
| Primer: kdpBC-comp-NdeI-F1:<br>5’ GCTAGG catatg GTG ATG ATC GCA CGC ATG GAG ACC T | This manuscript | N/A |
| Primer: kdpBC-comp-ClaI-R1:<br>5’ GCTACC atcgat TCA GAG GCG ATC CAA TGC GAG GTT | This manuscript | N/A |
| Primer: kstR1-nostart-BamHI-F3:<br>5’ GCTACC ggatcc TCG TCA GCG AAC ACG AAC ACC AGT | This manuscript | N/A |
| Primer: kstR1-HindIII-R1:<br>5’ GCTAGG aagctt CTA GGC GCT GTC TTG ATC GCC GAT CA | This manuscript | N/A |
